# GENESIS CGDYN: large-scale coarse-grained MD simulation with dynamic load balancing for heterogeneous biomolecular systems

**DOI:** 10.1101/2023.08.24.554724

**Authors:** Jaewoon Jung, Cheng Tan, Yuji Sugita

## Abstract

Residue-level coarse-grained (CG) molecular dynamics (MD) simulation is widely used to investigate slow biological processes that involve multiple proteins, nucleic acids, and their complexes. Biomolecules in a large simulation system are distributed non-uniformly, limiting computational efficiency with conventional methods. Here, we develop a hierarchical domain decomposition scheme with dynamic load balancing for heterogeneous biomolecular systems to keep computational efficiency even after drastic changes in particle distribution. The new schemes are applied to intrinsically disordered protein (IDP) droplet fusions. The droplet shape changes correlate with mixing IDP chains from two droplets. We also simulate formations of large IDP droplets, whose sizes are almost equivalent to those observed in microscopy. The methods have been implemented in CGDYN of the GENESIS software, which provides a new tool for investigating mesoscopic biological phenomena using the residue-level CG models.

## Introduction

Computational simulations with modeling of biomolecular structure and dynamics at various levels of detail can elucidate complex cellular phenomena in close collaboration with experimental studies. Quantum mechanics/molecular mechanics (QM/MM) or atomistic molecular dynamics (MD) methods provide detailed descriptions of the conformational dynamics of target molecular systems. However, they are computationally demanding when exploring long-time dynamics of a large biomolecule or a biomolecular system consisting of many biomolecules. Coarse-grained (CG) MD simplifies the description of these systems by representing multiple atoms as a single particle and thereby reduces computational complexity^1-3^. CG MD simulations retain essential structural and dynamic properties in biomolecules^3^, making them valuable for investigating biomolecular processes, such as protein folding and dynamics^4,5^, protein-DNA interactions^6,7^, nucleosome dynamics^8,9^, genome organizations^10,11^, condensate formation/destruction via liquid-liquid phase separation (LLPS)^12-14^, etc. Among many CG methods, residue-level CG MD simulations for proteins, nucleic acids, and lipids can bridge the gap between atomistic simulations and experimental observations^3^. They provide insights into fundamental but complex biological processes by balancing modeling accuracy and computational efficiency.

Various residue-level CG models have been developed to capture the structure, dynamics, and inter-molecular interactions of biomolecules. These models include the structure-based Go model^4^ and its variants for folded biomolecules (such as AICG2+^5^), the HPS model for intrinsically disordered proteins (IDPs)^13^, the 3SPN series models for nucleic acids^15-17^, and the Martini^18^ and SPICA^19^ models for lipid systems. Other notable CG models, for instance, UNRES^20^, OPEP^21^, and PRIMO/PRIMONA^22^ have been specifically designed to address different aspects of biological phenomena. Several MD programs, including Gromacs^23^, OpenMM^24^, NAMD^25^, HOOMD-blue^26^, LAMMPS^27^, Cafemol^28^, and GENESIS^29-31^, offer a diverse set of tools and capabilities for performing CG MD simulations in various contexts. While the CG models and MD programs have proven invaluable in addressing biological problems, there are difficulties in their implementations that do not arise in atomistic MD simulations. When CG MD simulation with the implicit solvent approximation is parallelized on multiple processors by the conventional domain decomposition scheme with an equal domain size for all processes, the processes in charge of dense regions have a significant workload. In contrast, those for dilute (or sparse) regions are almost idle, leading to non-negligible waiting time to synchronize tasks between all the processes. Moreover, the incorporation of diverse potential functions that describe interactions between different biomolecular components adds to the complexity of optimizing CG MD software^31^.

These difficulties limit the available system size even for residue-level CG models with the implicit solvent approximations. This requires innovative CG MD simulation schemes. Scientifically, there is a growing demand for residue-level CG MD simulations of large biological systems. For instance, protein/nucleic acid condensates (or droplets) formed by LLPS have attracted many chemists and biologists due to their relevance to serious neurotoxic diseases or essential biological functions in the cellular cytoplasm or nucleus^12, 13, 32, 33^. Currently, standard simulations of LLPS have used so-called slab models^13^, which have two short-length dimensions and only one long dimension within a periodic rectangular box. Dense and dilute phases are observed along the long dimension in equilibrium conditions. However, this anisotropic shape may not fully capture the three-dimensional (3D) nature of LLPS. Another example is the 3D modeling of chromatin. Many computational models of 3D structures of chromatin have been proposed using experimental data such as Hi-C^34, 35^. However, due to the dynamic nature of chromatin structures and computational limitations, most of those structural models are developed at relatively low resolutions (kilo-bases)^34, 35^. On the other side, at higher resolution, conformational dynamics of only a small number of nucleosomes with/without transcription factors were simulated with residue-level CG models^8, 9^.

Here, we develop a unique domain decomposition scheme with dynamic load-balancing to enable efficient residue-level CG MD simulations on parallel computers to handle non-uniform densities in a large biological system. Dynamic load-balancing is essential to accommodate rapid but significant changes in particle distributions in biological processes such as droplet formations from evenly distributed proteins. We have implemented the scheme in the GENESIS software^29-31^ as a new MD engine called “CGDYN” (CG molecular DYNamics), specifically designed for CG MD simulations. CGDYN has been optimized for parallel computers with many processors, outperforming other MD programs regarding computational performance. In this study, we utilize CGDYN to investigate molecular mechanisms underlying the fusion of two smaller droplets into a larger one using AICG2+^5^ and HPS^13^ models. Additionally, we conduct ultra-large-scale CG MD simulations consisting of multiple droplets to observe droplet formations, whose sizes are almost equivalent to the confocal microscope images. CGDYN in GENESIS provides a new computational tool for investigating mesoscale biological phenomena at the residue-level descriptions and connecting our understanding of the structure and dynamics of proteins and nucleic acids with cellular-scale biological phenomena.

## Results

### A new domain decomposition scheme with dynamic load balancing

We develop a novel domain decomposition scheme with dynamic load balancing to parallelize the residue-level CG MD simulations. The scheme is based on the midpoint cell method^36^ used in GENESIS SPDYN for atomistic MD simulations. The midpoint cell method divides a simulation space hierarchically: subdomains at first and then small cells from each subdomain. The cell size is decided from the interaction-range threshold, and the number of subdomains equals the number of processes. Because of almost uniform particle distributions in atomistic MD simulations, every subdomain has the same number of cells in SPDYN^29^. In contrast, each subdomain in CGDYN includes a different number of cells to avoid load imbalances by adopting a new domain decomposition scheme, which we call *the cell-based kd-tree method*. Here, subdomains for low particle density regions include more cells, while those for high-density regions contain fewer cells.

How to assign cells to each subdomain is described in Fig. 1a. We first divide a simulation system into two subdomains by making a boundary of cells such that two subdomains have nearly the same number of particles. Each subdomain is again divided into the next-level subdomains, in which their numbers of particles are almost identical. We iterate this procedure until the number of subdomains becomes the same as the process numbers. Each subdomain has particle data from the cell in the subdomain and the adjacent cells in other ones to compute bonded and nonbonded interactions (Fig. 1b). To complete this data structure, communication between subdomains, namely, sending particle coordinates from the boundary cells in one subdomain (i.e. the rank 5 in Fig. 1b) to the neighbors (the ranks 1, 4, 6, and 9) and receiving the coordinates from the boundary cells in the neighbors (the ranks 1, 4, 6, and 9) to the target subdomain (the rank 5), are required for the energy/force evaluation. In the residue-level CG MD simulations, particle densities in subdomains and cells can be changed rapidly. In such cases, the initial domain decomposition based on *the cell-based kd-tree method* cannot guarantee a good load balance. As an example, we show a simulation of the Heat-resistant obscure protein-11 (Hero-11)^37^ and the low-complexity domain of TDP-43 (TDP-43-LCD, with amino acid residues 261-414)^38^. TDP-43-LCD is known to form droplets in physiological conditions, while highly-charged Hero-11 can regulate the TDP-43-LCD condensate^37, 39^. At *t* = 0, the particle densities of the subdomains that include the TDP-43-LCD droplet are quite high, while the rest include just Hero-11 proteins at low concentrations. As the simulation progresses, the particle densities become more uniform, requiring different domain decompositions from the initial one (Fig. 1c). During the simulation, we decompose the simulation space using the *cell-based kd-tree method* at a fixed interval (around 10^6^ steps but it depends on the integration time step and the target system). This example suggests the importance of dynamic load balancing to keep the best performance of long-time CG MD simulations.

**Figure 1.**
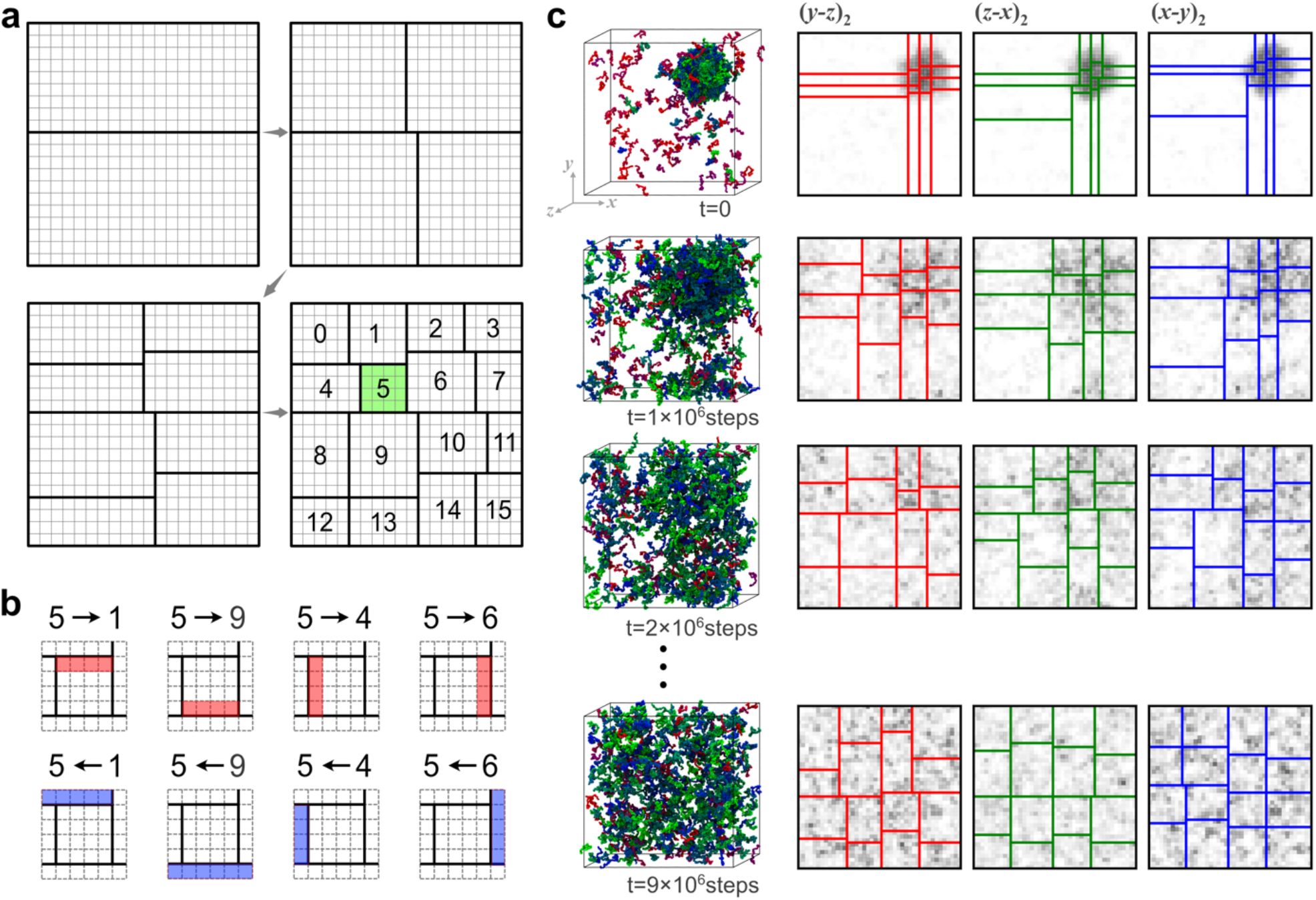
(a) The domain decomposition algorithm with the cell-based kd-tree method. (b) Communication in sending the coordinates from MPI rank 5 to other processes (upper) and receiving the coordinates from other processes to MPI rank 5 (lower). (c) Updates of subdomains during MD simulations of Hero11 and TDP-43-LCD.

### Benchmark tests of CGDYN for heterogeneous biological systems

We first examine the computational performance of CGDYN in CG MD simulations of heterogeneous biological systems. Two multiple-droplet systems, whose particle densities are different, are prepared for the performance comparison: one with *ρ* =1.85×10^−6^ and the other with *ρ* = 4.97 ×10^−7^ (chains per Å^3^, Fig. 2a). Both systems consist of 1,949 TDP-43-LCD chains (number of particles is 300,146). We compared the computational performances for three different algorithms in GENESIS: (i) CGDYN (the cell-based *kd-*tree method with dynamic load balancing), (ii) SPDYN-like (the original mid-point cell method without dynamic load balancing), and (iii) ATDYN (atomic decomposition without dynamic load balancing). ATDYN shows no performance dependence on the two systems, but the computational efficiency is limited to 2.0 ×10^6^ steps/day (Fig. 2b). In comparison, MD simulations using CGDYN show 3∼30 times better performances than ATDYN (Fig. 2b). Importantly, almost identical speeds are obtained between the high and low-density systems with CGDYN due to the efficient parallelization with dynamic load balancing, suggesting that these algorithms used in CGDYN work well irrespective to the particle densities. MD simulations with CGDYN accelerate at most 7.5 times compared to the SPDYN-like algorithms (Fig. 2b). The acceleration with CGDYN can depend on the frequency of the dynamic load balancing. Supporting Fig. 1 shows that the twice better performance is observed by applying the dynamic load balancing 100 times more frequently.

**Figure 2.**
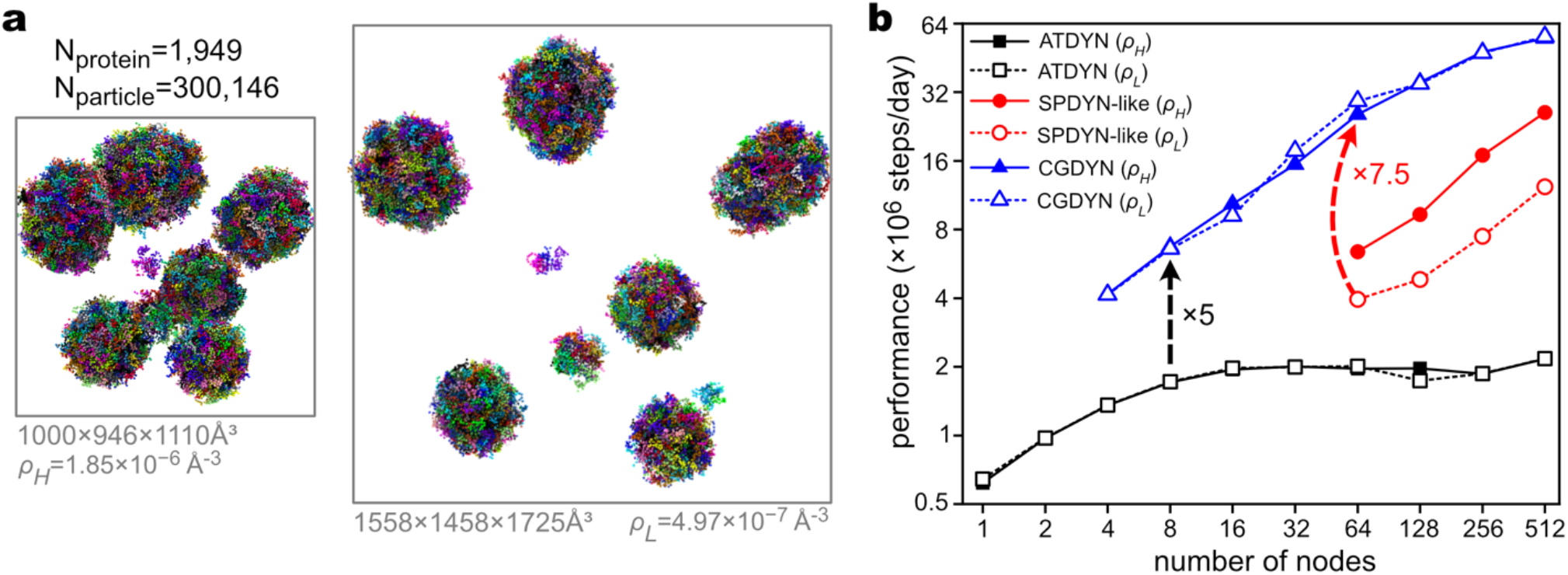
Benchmark results of CG MD simulations with three algorithms: ATDYN, SPDYN-like and CGDYN. (a) The two benchmark systems with different densities. (b) Benchmark performance (× 10^6^ steps/day). ATDYN can be used with a small number of nodes, but the parallel efficiency is low, and the performance is saturated from 16 nodes. The SPDYN-like algorithm performs better using many nodes but has a higher dependency on particle density. CGDYN performs better than MD simulations based on the ATDYN and SPDYN-like algorithms, showing a weak dependency on particle densities.

We find that CGDYN works well for very large systems with more than 2.5 million CG particles (Supporting Fig. 2). It shows good scalability up to 4,096 nodes on Fugaku (16,384 MPI processes) with ∼5.0 ×10^7^steps/day, promising to simulate very large biological systems efficiently with residue-level CG models. Supporting Fig. 3 shows the performance comparison of residue-level CG MD simulations of DNA systems between ATDYN, CGDYN, and Open3SPN2^40^. Although Open3SPN2 is based on OpenMM^24^ and accelerated with GPU processors, CGDYN on a single CPU performs better on multiple double-stranded DNA systems than Open3SPN2. For larger DNA systems, the superiority of computational performance with CGDYN is more significant, promising efficient residue-level CG MD simulations of mesoscopic biological systems (Supporting Table 1).

We also perform benchmark tests for the systems already published in our previous publication. CGDYN has memory problems on the RIKEN supercomputer Hokusai, using a small number of processes for large systems (512 nucleosomes). However, we could run MD even for very large systems by increasing the number of processes, and finally, CGDYN outperforms ATDYN on all systems with more than ten-fold speed up (Supporting Fig. 4).

### Molecular mechanisms for the fusion of two droplets

The fusion of phase-separated liquid-like droplets is an important mechanism for maintaining stable cellular environments when the concentration of components changes^41^. The residue-level CG MD simulations^12, 13, 33^ allow the investigation of inter-molecular interactions and the resulting diffusion/mixing of different components during the droplet fusions. However, MD simulations of this process are computationally demanding and not frequently employed in practical studies due to the large number of molecules involved and the rapid exchange of components in droplets. In this study, we use CGDYN to examine the fusion of two separated droplets, each consisting of approximately 500 chains of TDP-43-LCD (resulting in 1,000 chains in total, Fig. 3a). The HPS model^13^ was utilized for the entire protein, except for a short α-helical region (residues 320 to 334) modeled with AICG2+^5^.

**Figure 3.**
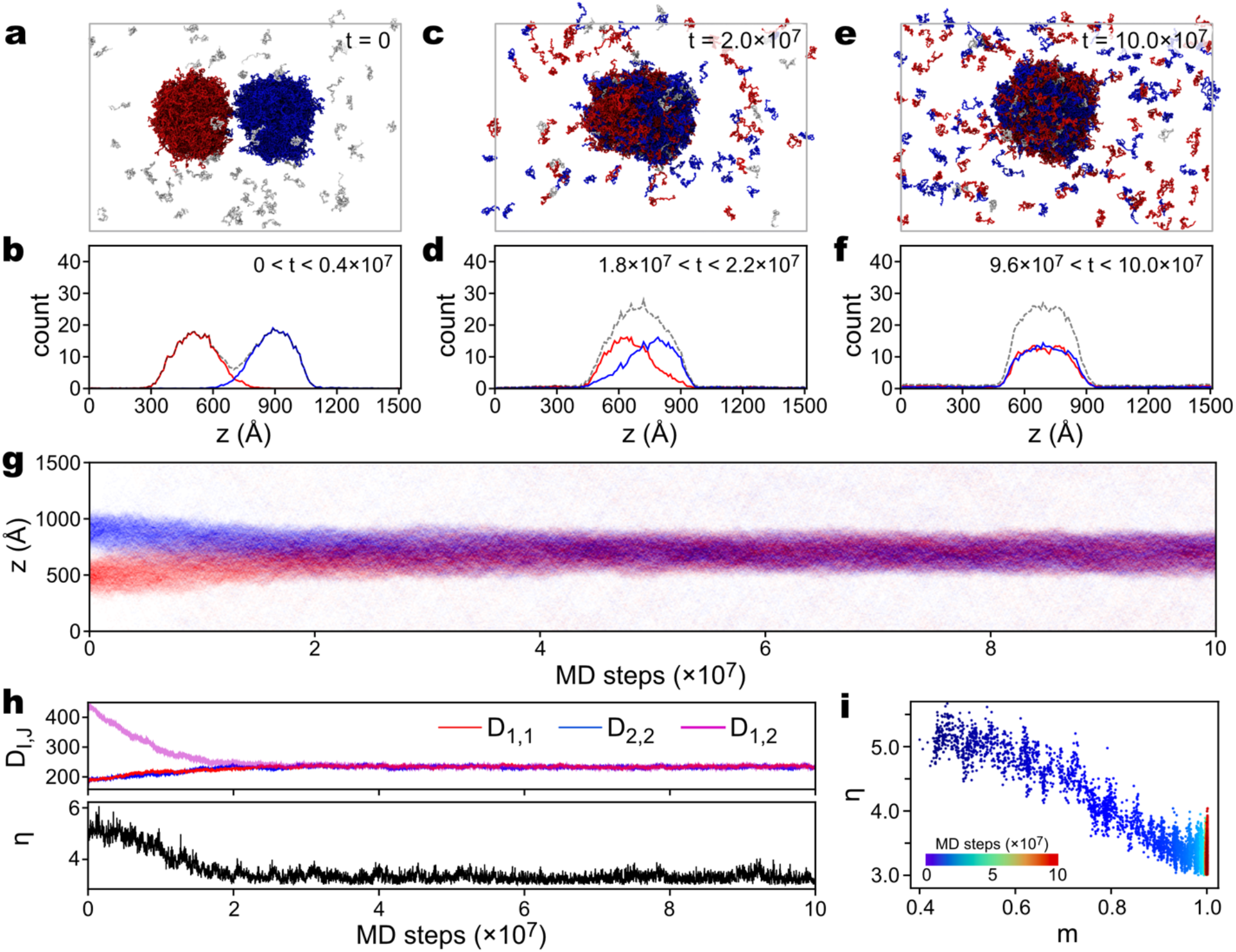
CG MD simulation of the fusion process of two TDP-43-LCD droplets. The system consists of 1,000 chains of TDP-43-LCD. In the initial structure, two separate droplets were put close to each other. The system was simulated at 280K for 10^8^ steps. (a, c, and e) Snapshots of simulated structures at *t* =0 (a), *t* =2 ×10^7^ steps (c), and 1 ×10^8^ steps (e). Chains from the two droplets in the initial structure are colored as red or blue, respectively. (b, d, and f) Time-averaged distributions of TDP-43 chains along the *z* axis during 0 < *t* < 0.4 ×10^7^ (b), 1.8 ×10^7^ < *t* < 2.2 ×10^7^ (d), and 9.6 ×10^7^ < *t* < 10.0 ×10^7^ (f) steps, respectively. (g) Density of TDP-43 particles along the *z* axis as a function of simulation time. (h) Time series of average chain-chain distances *D*_*I,J*_ (upper) and shape coordinate *η*(lower). (i) Droplet mixing (depicted by coordinate *m*) against shape change (*η*).

The MD simulations were conducted at *T* =280*K* for 1 ×10^8^ steps to explore the dynamics of the system. Despite rapid density changes during the simulations, we achieved a speed of 1.25 ×10^7^ steps per day on a single node (16 MPI processes in conjunction with 3 OpenMP threads) on a local PC cluster (Intel Xeon Gold 6240R CPU, 2.4GHz). The DBSCAN clustering analysis^42^ is employed to identify chains belonging to the two distinct droplets in the initial structure (Figs. 3a and 3b). By labeling each chain, we can track their positions and monitor the fusion process (Figs. 3a to 3g). Interestingly, the fusion of two droplets occurs shortly after the start of the MD simulation and completes within the first several 10^7^ MD steps (Fig. 3g). To investigate the interaction and redistribution of components within the droplets during fusion, we monitored the “mixing” of contents and the associated shape changes.

Through structural analysis and calculation of chain distribution in the simulation box, we observed that the two droplets merge into one at approximately 2 ×10^7^ steps, although the shape of the merged droplet is a flattened ellipsoid, and the chains from the original droplets are not yet fully mixed (Fig. 3c and 3d). After 1 ×10^8^ steps of simulations, the chains from both droplets mixed completely (Figs. 3e and 3f). To quantify the relationship between the content mixings and the shape changes, we introduce five order parameters. One parameter, *D*_*I,J*_, represents the average distances between the center-of-masses (COMs) of chains in the original droplets *I* and *J* (*I, J* =1 or 2, see Methods). *D*_1,1_ and *D*_2,2_ increase during the mixing process, while *D*_1,2_ decreases (Fig. 3h), depicting the redistribution of chains from their original droplet into the fused one. We further define *m* as 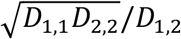 to incorporate information from *D*_1,1_, *D*_2,2_, and *D*_1,2_. As expected, *m* increases during the fusion process. Another coordinate, *η*, is used to describe the anisotropy of the droplet during fusion (see Methods for definition). Notably, we observe that mixing (*m*) and shape change (*η*) are coupled, with the fused droplet reaching a sphere-like shape (*η*< 4.0) slightly earlier than the complete mixing of contents (*m* ≈ 1.0, Fig. 3i).

### Toward the observation of ultra-large droplets that are detectable by experimental confocal microscopy

In contrast to the droplet formation or regulations studied in MD simulations, the real droplet sizes observed with confocal microscopy vary from sub-micrometer to tens of micrometers^43, 44^. To connect a large gap between the simulations and experiments, we focus on the fusion of multiple droplets toward a much larger one, which happens in the cell during the formation of mesoscopic-scale protein droplets due to LLPS^43, 45, 46^. Here, we generated large systems consisting of 16,657 chains of TDP-43-LCD, resulting in 2,565,178 CG particles. Two systems were created, each with different overall chain densities: 1.85×10^−6^ Å^-3^ and 6.75 ×10^−7^Å^-3^, respectively (Figs. 4a and 4b). The high- and low-density systems were simulated in boxes measuring 2087 ×2067 ×2077 Å^3^ and 2899 ×2903 ×2929 Å^3^, respectively. Both simulations were performed in the NVT ensemble at *T* =290*K*. Leveraging the computational power of the supercomputer Fugaku, we achieved a simulation speed of 2.8 ×10^7^ steps per day (∼280 ns/day, with a time integration step size of 10 fs) on 512 nodes, irrespective of the particle densities of the two systems.

**Figure 4.**
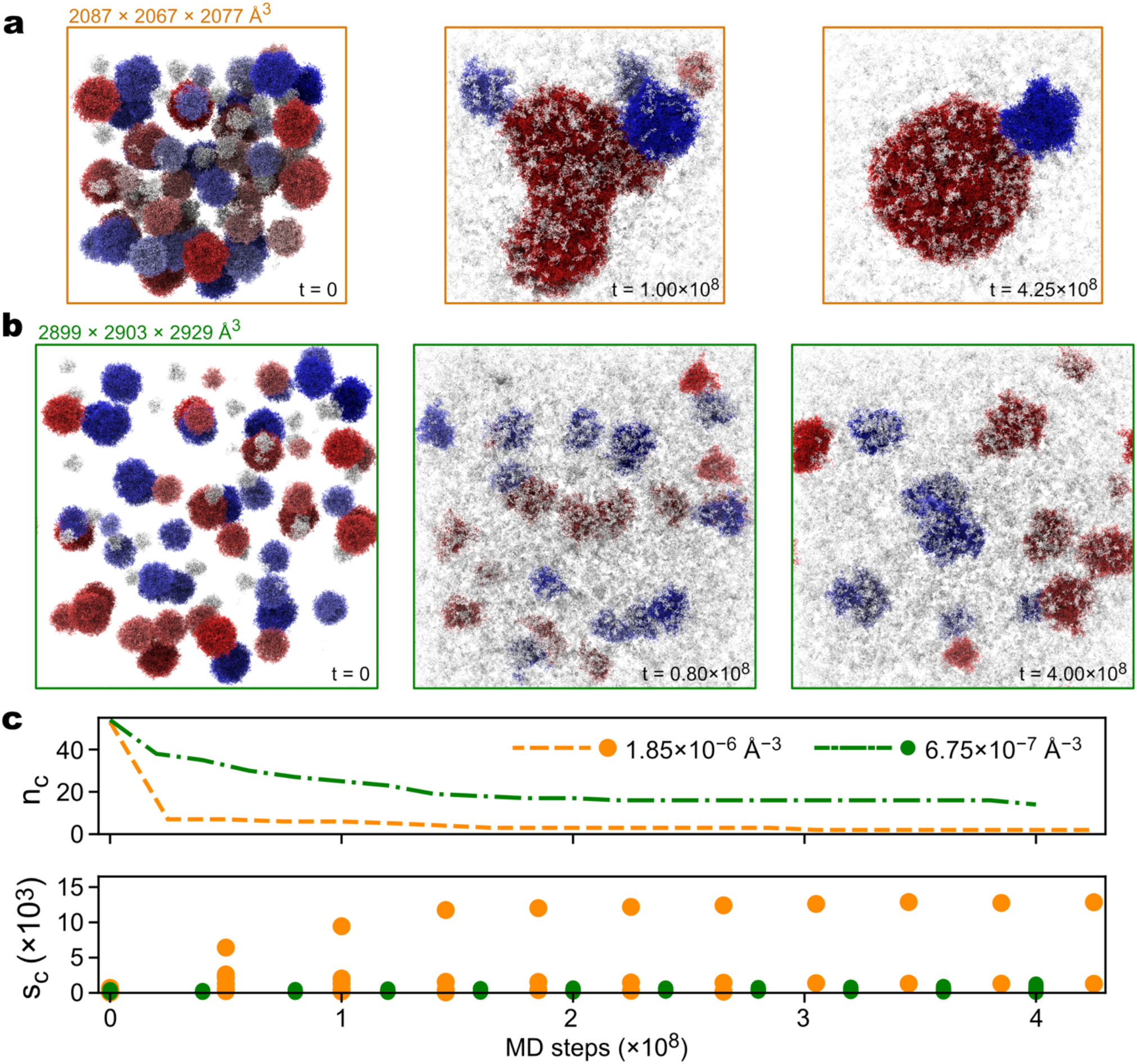
CG MD simulations of multiple TDP-43-LCD droplet fusions at two different densities. (a) Snapshots of the high-density system taken at three different times during the simulation. (b) is the same as (a) but for the low-density system. (c) Number (*n*_*c*_, upper) and sizes of clusters (*s*_*c*_, lower) as functions of simulation time.

To monitor the fusion of small droplets into larger ones, we employ the DBSCAN clustering analysis^42^ on selected snapshots obtained at different stages during the simulations. In the initial structures, proteins are distributed into more than 50 droplets of varying sizes, ranging from 5 to 452 chains in a droplet. While the two systems exhibit similar cluster size distributions in the initial structures, more drastic changes happen in the high-density system as the simulation progresses. In the high-density system, the number of clusters (*n*_*c*_) quickly decreases from *n*_*c*_ =53 to *n*_*c*_ =7 within the first 5 ×10^7^ steps, while the largest cluster size increases from *s*_*c*_ =452 at *t* =0 to *s*_*c*_ =6422 at *t* =5 ×10^7^ steps (Fig. 4c), indicating the occurrence of droplet fusions. Indeed, a snapshot at 1.00 ×10^8^ steps of the high-density system reveals the formations of a few large clusters of TDP-43-LCD, with the largest cluster exhibiting a non-spherical shape (Fig. 4a). After 4.25 ×10^8^ steps, the seven clusters further assembled into two droplets, with the larger droplet comprising 12,842 chains and a diameter of approximately 0.1 μm (Fig. 4a). In contrast, the number of clusters in the low-density system decreases at a slower pace, reducing from *n*_*c*_ =53 to *n*_*c*_ =35 during the first 4 ×10^7^ steps and eventually reaching 14 after 4 ×10^8^ steps (Fig. 4c). Correspondingly, the largest cluster size only increases from *s*_*c*_ =452 (*t* =0) to *s*_*c*_ =491 at *t* =2 ×10^7^ steps and finally to s_+_ =1,261 at *t* =4 ×10^8^ steps. These results demonstrate the dependence of fusion behavior on the overall concentration of IDPs.

These simulation results highlight the effectiveness of CGDYN in capturing the fundamental events involved in droplet fusions with residue-level particle resolutions. These initial results underscore the potential for gaining valuable insights into the fusion mechanisms. We expect that further extensive simulations and in-depth analysis can uncover additional details and refine our understanding of the IDP droplet fusion processes.

## Discussion

Slab simulations have been commonly employed to investigate equilibrated phase behavior of biomolecular condensates^12, 13, 33, 38, 39^. It is crucial to note that these simulations have limitations in accurately capturing the 3D nature of the fusion processes due to the limited number of chains and the small size of the system. In contrast, the large-scale droplet simulations with the residue-level description of proteins can overcome these potential problems in smaller-scale simulations. As demonstrated in the current study, CGDYN is a powerful tool for simulating realistic-sized biomolecular condensates. The versatility of CGDYN could facilitate the reproduction of many experimental results, such as fluorescence recovery after photobleaching^38^ and optical tweezers^47^, and sheds more light on the molecular mechanisms underlying the biomolecular condensates.

The LLPS of biomolecules and the subsequent transitions from liquid to solid phases can be influenced by specific proteins^37^ or RNA molecules^48, 49^. In a recent study, we explored the regulation of TDP-43-LCD’s condensation by Hero-11^39^. Through slab simulations simulated with ATDYN, we could investigate the effects of Hero-11 on the interactions and dynamics of TDP-43-LCD, including its potential influence on droplet fusion through charge distribution. We expect that by incorporating Hero-11 into the TDP-43-LCD droplet systems examined in this work, we can provide direct evidence for one of the previously proposed mechanisms and develop a deeper understanding of the regulation of biomolecular LLPS.

The internal structures of droplets or mesoscopic assemblies composed of multicomponent proteins/RNAs have attracted significant attention^50-54^. Computational and experimental studies have focused on investigating the multi-layered structures in multi-component systems^50-52^. Valency has been proposed as a key factor in determining the spatial positioning of each component^52^. The impact of layered distributions, such as reducing the surface tension and localizing higher-valency components at the core while allowing rapid exchange of lower-valency components in and out of condensate, has been discussed^52^. The dynamics of multi-component LLPS has been proposed to involve multiple phase transitions, including the first separation of condensates from the bulk solution and subsequent transitions within the high-density phase^55^.

There are increasingly interesting mesoscopic biological phenomena to be simulated with large-scale CG MD simulations with residue-level CG models. Paraspeckles, which are condensates composed of RNAs and IDPs, are observed in the cellular nucleus of mammalian cells^56^. A triblock copolymer model has been proposed to explain the shell localization of RNA ends and the size of paraspeckles^57^. However, due to their large sizes, achieving residue-level descriptions of such biologically interesting systems has been challenging. Multi-dimensional information on chromatin is also accumulated, providing a deeper understanding of how gene expression is regulated via dynamic interactions involving nucleosomes, transcription factors, remodelers, RNA polymerases, and other factors^58-61^. Experimental studies have suggested the functional roles of liquid droplets formed by transcription factors, mediators, and RNA polymerases^59-61^. To understand these models structurally and epigenetically, the residue-level descriptions are necessary at the very least. The methods and software developed in this study could be important computational bases for understanding mesoscopic biological phenomena through long-time dynamics of the real-size simulation systems.

## Methods

### Potential functions of the residue-level CG models

In GENESIS CGDYN, we employ residue-level CG models with an approximate resolution of 10 heavy atoms per particle. At this level, each CG particle represents a single amino acid residue in proteins, while nucleic acids are represented by three particles per nucleotide, corresponding to the phosphate (P), sugar (S), and base (B) components.

### Protein models

Regarding proteins, we have incorporated two distinct CG models: one for folded domains (AICG2+^5^) and another for IDRs (HPS/KH^13^). These models can be incorporated for proteins comprising folded domains and IDRs.

The AICG2+ potential energy function is given by^5^:

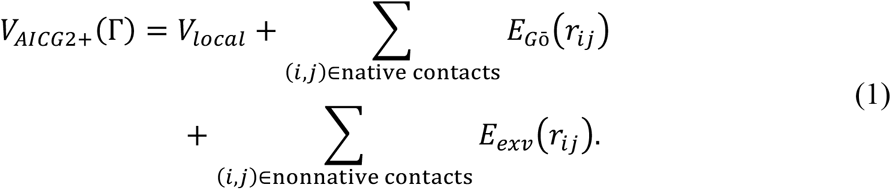

where Γis the conformation of protein, *V*_*local*_ includes all bonded terms, *E*_*G*ō_(*r*_*ij*_) is the structure-based Gō potential, and *E*_*exv*_ (*r*_*ij*_) is the potential from excluded volume interaction.

The local interaction term, *V*_*local*_ in Equation (1), is defined as^5^:

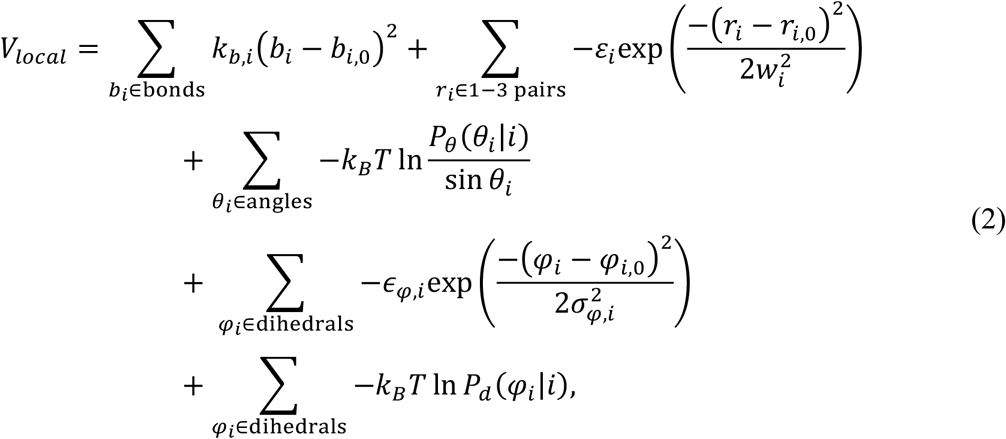

where the first term is for the bond interaction, the second term is for every two end particles in the angle one, and the fourth term is for the dihedral angle potential. The third and fifth terms are statistical flexible potentials, where *P*_*θ*_(*θ*_*i*_|*i*) and *P*_*d*_(*φ*_*i*_|*i*) are residue-type dependent probability distributions of angles and dihedral angles, respectively. *k*_*B*_ is the Boltzmann constant and *T* is temperature.

*E*_*G*ō_ *r*_*ij*_ in Equation (1) is given by:

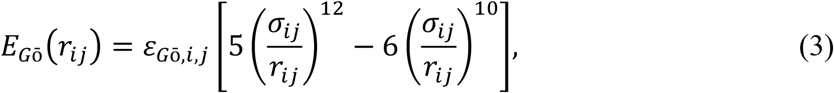

where *r*_*ij*_ is the distance between residues forming a native contact, *σ*_*ij*_ is the reference value of *r*_*ij*_, and *𝜀*_*G*ō*i,j*_ is the context-dependent energy coefficient. A native contact is defined as two residues having any heavy atoms within 4.5Å from each other in the reference structure. Only the folded regions of proteins have *E*_*G*ō_(*r*_*ij*_) interactions.

*E*_*exv*_(*r*_*ij*_) in Equation (1) is given by:

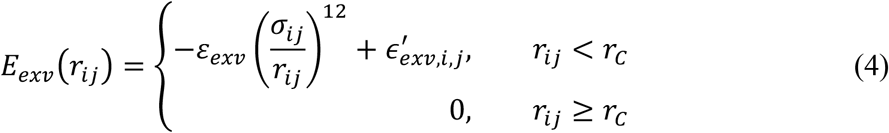

where *r*_*ij*_ is the distance between residues *i* and *j, σ*_*ij*_ is residue-type dependent excluded volume distance^6^, 𝜀_*exv*_ =0.6*kcal*/*mol* is the force coefficient, *r*_*C*_=2*σ*_*ij*_ is the cutoff distance, and 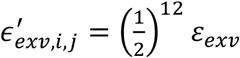.

The potential energy function of the HPS model is defined as^13^:

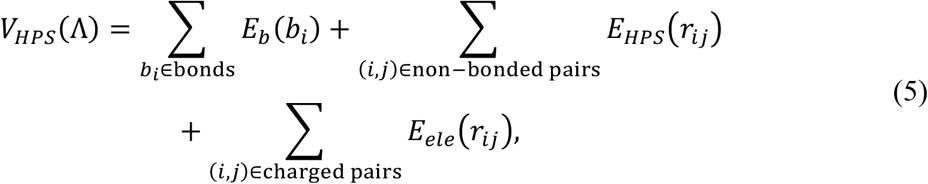

where Λ is the conformation of an IDP and *E*_*b*_(*b*_*i*_) is the potential for every two neighboring particles with a bond length *b*_*i*_:

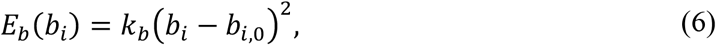

where *b*_*i*,0_ =3.8Å is the reference value and *k*_*b*_ =2.39 *kcal*/*mol* · Å^-2^ is the force constant.

*E*_*HPS*_(*r*_*ij*)_ is the interaction between non-bonded particles^13^:

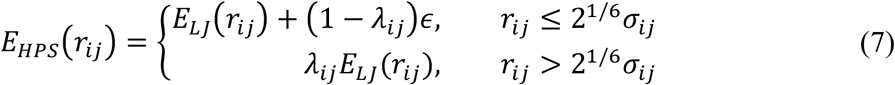

where *λ*_*ij*_ is the hydropathy and *E*_*LJ*_(*r*_*ij*_) is the Lennard−Jones potential:

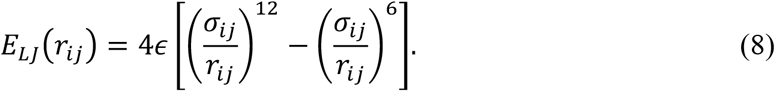

In Equations (7) and (8), *𝜖* =0.2*kcal*/*mol*. We use an arithmetic combinational rule for *σ*_*ij*_ and *λ*_*ij*_.

We utilize the Debye-Hückel term for the electrostatic interaction *E*_*ele*_(*r*_*ij*_):

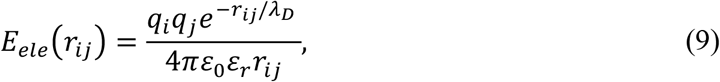

where *r*_*ij*_ is the distance between two non-bonded charged particles *i* and *j* and 𝜀_0_ is the dielectric permittivity of vacuum. 𝜀_*r*_, the relative permittivity of the solution, is defined as a function of the temperature *T* and salt molarity *C* : 𝜀_*r*_ =*e*(*T*)*a*(*C*), where *e*(*T*) =249.4 − 0.788 *T* + 7.20 ×10^-4^*T*^2 62^ and *a*(*C*) =1 − 0.2551*C* + 5.151 ×10^-2^*C*^2^ − 6.889 ×10^-3^*C*^3 63^. The Debye length *λ*_*D*_ is given by 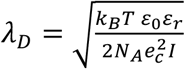, where *e*_*c*_ is the elementary charge, *N*_*A*_ is the Avogadro’s number, and *I* is the ionic strength of the solution.

### Nucleic acid models

We utilize the 3SPN.2C model for DNA, whose potential energy function is defined as^15, 16^:

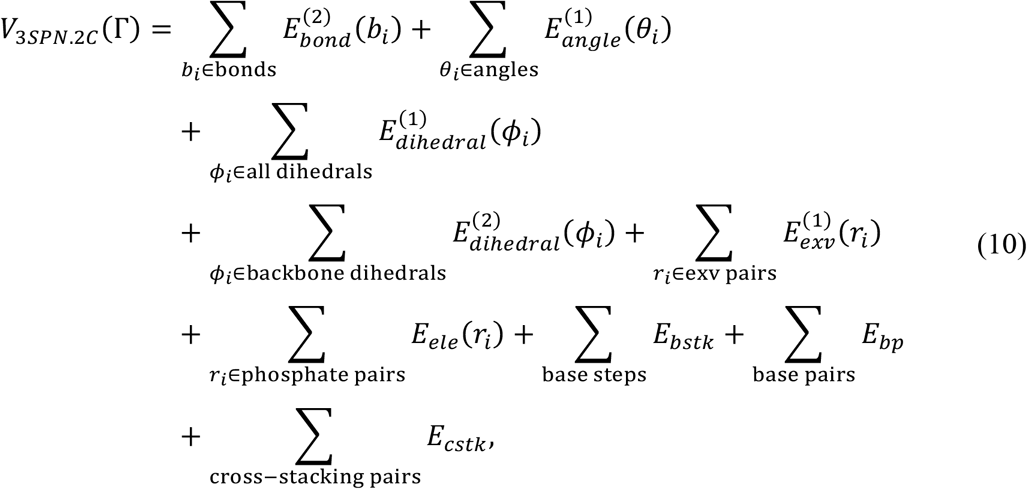

where Γ represents the conformation of DNA. Here, “excluded volume pairs” (shown as exv pairs in eq. (10)) are the nonbonded particle pairs that do not participate in base-pairing or stacking interactions.

Bond potential in Equation (10) is defined as^15, 16^:

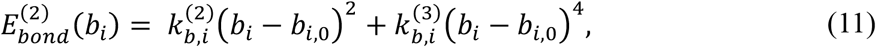

where *b*_*i*_ is the bond length, *b*_*i*,0_ is the reference value of *b*_*i*_, 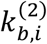 and 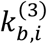 are the force constants in the quadratic and quartic terms, respectively.

The angle potential in Equation (10) is given by^15, 16^:

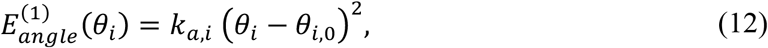

where *θ*_*i*_ is the bond angle formed by three CG particles, *θ*_*i*,0_ is the reference value, and *k*_*a,i*_ is the force constant.

The dihedral angle potentials in Equation (10) are defined as^15, 16^:

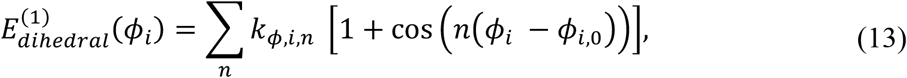

and:

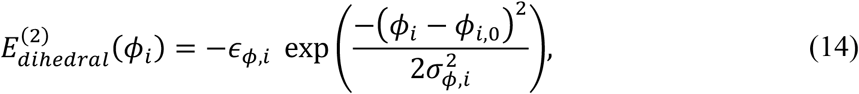

where *ϕ*_*i*_ and *ϕ*_*i*,0_ are the dihedral angle and its reference value, respectively, *n* is an integer number that controls the periodicity of the function, *σ*_*ϕ,i*_ is the Gaussian width, and *k*_*ϕ,i,n*_ and *𝜖*_*ϕ,i*_ are the force constants.

The terms *E*_*bstk*_, *E*_*bp*_, and *E*_*cstk*_ in Equation (10) refer to multi-body energy functions describing base-base interactions^15, 16^:

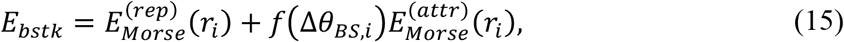

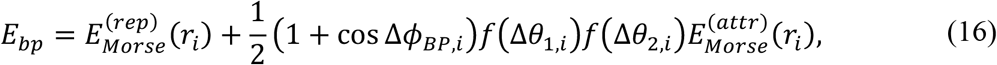

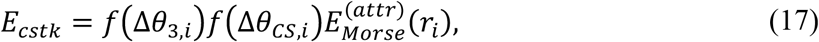

where *r*_*i*_ represents the distance between the two interacting bases, and the angles (*θ*_*BS,i*_, *θ*_1,*i*_, *θ*_2,*i*_, *θ*_3,*i*_, and *θ*_*CS,i*_) and dihedral angles (*ϕ*_*BP,i*_) are formed by the surrounding sugar and phosphate sites. The Morse potential in Equations (15), (16), and (17) are defined as:

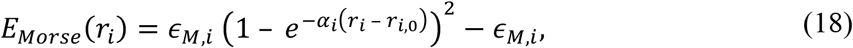

where *r*_*i*_ and *r*_*i*,0_ are the distance between two particles and its reference value, respectively. *ϵ*_*M,i*_ and *α*_*i*_ are the “depth” and the “width” of the Morse potentials, respectively. The repulsive 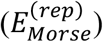 and the attractive 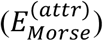 components of the Morse potential are defined in the following:

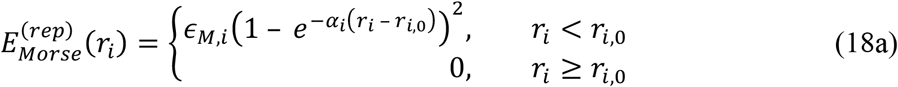

and

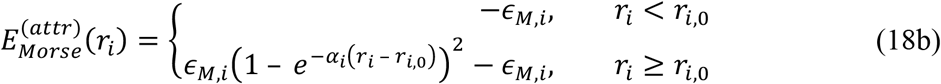

The angle-dependent modulating function in Equations (15), (16), and (17) is defined as^15, 16^:

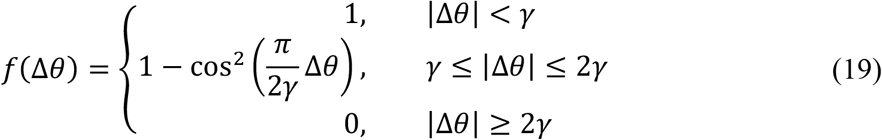

where Δ*θ* is the difference between an angle (*θ*) and its reference value, and γ controls the tuning range.

The excluded volume interaction in Equation (10) is given by^15, 16^:

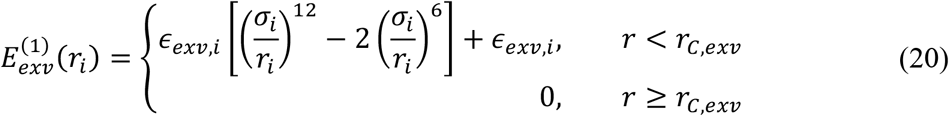

where *r*_*C,exv*_ is the cutoff distance and has the same value as *σ*_*i*_.

For the electrostatic interactions in Equation (10), we use the same definition given by the Debye-Hückel model as in Equation (9).

For RNA, we offer two models: a structure-based model (similar to Equation (1)) with a three-bead-per-nucleotide resolution^64^, and an HPS model (Equation (5)) with a one-bead-per-nucleotide resolution^33^.

### Protein-DNA Models

Protein-DNA binding can be divided into two types: sequence-nonspecific interactions involving amino acids and DNA backbone groups (primarily electrostatic interactions), and sequence-specific interactions between amino acids and DNA bases. The PWMcos model can describe the latter by incorporating position weight matrix (PWM) information into structure-based interactions^65^. This model identifies a set of DNA-binding protein residues (DB-*C*_*α*_s) that contact DNA in its native structure. The potential energy is then calculated using:

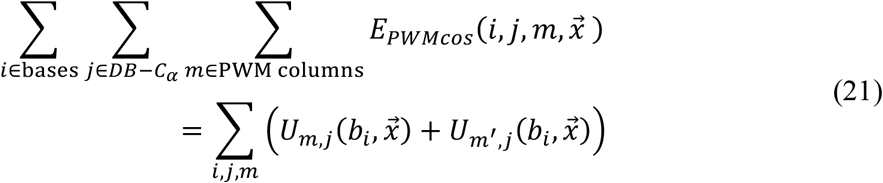

where *m*′ is the base in the complementary base of *m, b*_*i*_ is the base type of base *i* (*b*_*i*_ ∈ [*A, C, G, T*]), and 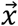 is the coordinates of particles in each conformation.

Function *U*_*m,j*_(*b*_*i*_, 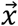) is defined as^65^:

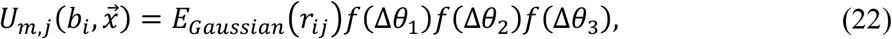

where *r*_*ij*_ is the distance between the *i*-th base and the *j*-th *C*_*α*_, and *θ*_1_, *θ*_2_, and *θ*_3_ are angles defined by the surrounding particles^65^.

The Gaussian potential is defined as^65^:

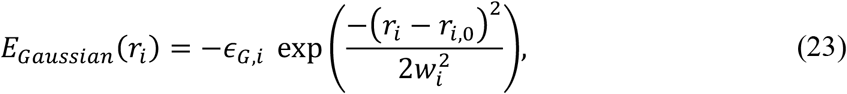

where *ϵ*_*G,i*_ and *w*_*i*_ are the “depth” and “width” of the Gaussian, respectively. *𝜖*_*G,i*_ is a constant dependent on base type and the PWM. The modulating functions *f*(Δ*θ*) is defined in Equation (19).

Similar to PWMcos, we also implement a sequence-nonspecific model to describe the hydrogen bond (HB) interactions formed between protein and DNA backbone^9^.

### CGDYN structure

In CGDYN, we implemented *the cell-based kd-tree scheme* as a domain decomposition scheme for heterogeneous biological systems. The cell size is greater than or equal to half of the distance wherein the neighbor list for electrostatic interaction is considered. All particle information in each subdomain (coordinate, force, charge, atom class number, and so on) is saved cell-wise (Supporting Fig. 5). We make an identifier of particles (ordered/disorder protein region, DNA base, and so on) for efficient neighbor list generation. For each cell, we first locate the information on charged particles followed by uncharged particles in this order. The array of potential function type and parameters of bonded interactions (bond/angle/dihedral angle list) is prepared without using cell indices (Supporting Fig. 5). In the case of bond, the program writes parameters of quadratic and next quartic bond terms sequentially in bond-related array. There are three potential functions for angle terms according to Equations (2) and (12):

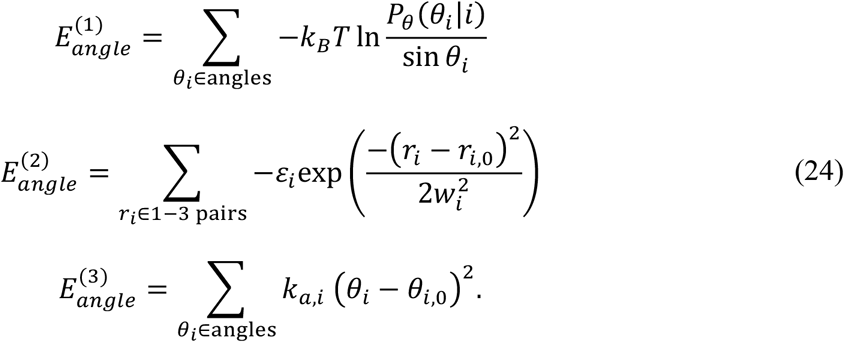

The program first writes parameters of 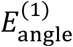 and next writes parameters 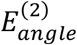 and 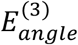 .

Similarly, we have three dihedral angle potential functions:

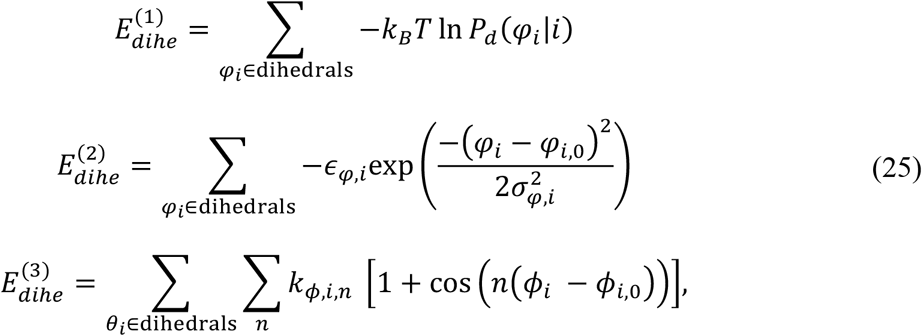

and the program writes parameters sequentially in dihedral angle array.

All nonbonded interactions, including bonded, electrostatic, HPS, excluded volume, etc, have different interaction ranges. Among them, electrostatic interaction has the longest interaction range. For a given particle, the interaction range and the cell involved in the calculation are considered as Supporting Fig. 6. For bonded and excluded volume interactions, we consider the particles in the target cell and neighboring cells. Electrostatic and HPS potential functions have longer interaction ranges. Therefore, particles in the next neighboring cells from the target cell are considered.

We generate the neighbor lists concurrently for all nonbonded interactions except for electrostatic and protein-DNA terms (Supporting Algorithm). First, we predefine threshold values of the neighbor list generation for excluded volume, DNA pairing, and HPS, which are denoted here as *r*_p,exv_, *r*_p,dna_, *r*_p,hps_, respectively. If a pairwise distance between two particles is shorter than *r*_p,exv_, we consider the particles in the neighbor lists for all the nonbonded interactions. If the distance, *r*, is in the range of *r*_p,exv_< *r* < *r*_p,dna_, we do not consider the particles in the neighboring list of excluded volume. If the distance exceeds *r*_p,hps_, the particles are included in the neighbor lists of the HPS interactions. The neighbor lists of electrostatic interaction are generated separately by evaluating pairwise distances only between charged particles with predefined threshold value, *r*_p,ele_. Threshold values of these interactions in the neighbor search are greater than those in the nonbonded energy/force calculations with *r*_p,XX_ =*r*_c,XX_ + *r*_buffer_ (*r*_p,XX_ and *r*_c,XX_ are threshold of neighbor list generation and energy/force calculations for XX interactions, respectively). Neighbor lists for PWMcos/HB potential used in protein-DNA interactions are also considered separately. The frequency of neighbor list search is user-defined value, but the program skips the neighbor search if the maximum particle displacement is less than half of *r*_buffer_.

To accelerate the evaluations of nonbonded interactions, SIMD (Single Instruction, Multiple Data) is applied to electrostatic and HPS force calculations by doing calculations even for unnecessary pairs with the pairwise distance between *r*_c,XX_ and *r*_p,XX_ . For other interactions including the excluded volume, DNA base pairing, SIMD is not applied because the ratio of 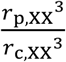 is much larger than 1 and it is better to skip unnecessary calculations for the pairwise distance range between *r*_c,XX_ and *r*_p,XX_ using conditional statements instead of applying SIMD.

Dynamic load balancing with the cell-based kd-tree scheme is applied at fixed intervals (which we will name the load balance update period), defined by application users. In this procedure, we do not change the cell size and only change the cell assignment to each subdomain. This process is applied as a default when we start MD. If the MD simulation time is the multiple of the load balance update period, the cell reassignment to each subdomain is done in a procedure like Fig. 1a. Communication between neighboring processes is redefined in a procedure like Fig. 1b. The subdomain information is saved in the global data array and the global data is completed by collective communication. Each subdomain reads new subdomain information from the global data and MD simulations continue based on the new subdomain data (Supporting Fig. 7).

### Modeling of TDP-43-LCD

We used the AlphaFold2^66^ predicted structure of TDP-43 as a reference structure for the residue-level CG modeling. In this modeling approach, each amino acid was represented by a single CG particle. We employed the HPS model^13^ for simulating TDP-43-LCD (residues 261-319 and 335 to 414), mostly as an intrinsically disordered protein (IDP). To preserve the secondary structure of an α-helix (residues 320 to 334), we employed the AICG2+ model^5^. The GENESIS-CG-tool is used to prepare all the structure and topology files for the MD simulations^31^.

### Preparation of initial structures for the droplet systems

We constructed the initial structures for the droplet simulations in a stepwise manner. First, we simulated a single chain of TDP-43-LCD for 2 ×10^7^ steps. From this simulation trajectory, we randomly selected a structure, which was then duplicated to create a system consisting of 500 chains. Subsequently, we performed “shrinking” simulations, gradually compressing the simulation boxes to dimensions of 500Å ×500Å ×500Å. The system was equilibrated at 260K, resulting in the formation of a single droplet of TDP-43-LCD. Next, we selected structures obtained from the single-droplet simulations and put them into a larger simulation box to construct multiple-droplet systems.

Specifically, we constructed the following systems:

1. We constructed a system consisting of 500 TDP-43-LCD chains and 100 Hero11 chains (Fig. 1c). We selected one simulated structure of the single TDP-43-LCD droplet system and placed it randomly within a simulation box of 1000Å ×1000Å ×1000Å. 100 Hero11 chains were then randomly added to the empty space, ensuring there were no structure clashes.
2. To construct the multiple-droplet systems in Figs. 2 and 4, we first carried out MD simulations of meso-scale particles with radii ranging from 50 Å to 200 Å. Only excluded volume interactions were considered during these simulations. Various numbers of large particles were simulated within simulation boxes of different sizes, to achieve varying densities. Next, we superimposed the single-droplet TDP-43-LCD structures onto the large particles and removed any TDP-43-LCD chains that located beyond the boundaries of the large particles. This process allows us to obtain multiple-droplet systems without structure conflicts.
3. To construct the initial structures of the two droplet simulations (Fig. 3), we chose two structures (named Λ_1_ and Λ_2_) from the single-droplet simulations of TDP-43-LCD. These two selected structures were placed into a simulation box with dimensions of 1000Å ×1000Å ×1500Å, ensuring that the two droplets in each structure were positioned close to each other with no direct contact (Fig. 3a). In the merged structure, certain chains in the dilute phase of Λ_1_ overlapped with the condensate in Λ_2_. For these chains, we relocated them to random locations in the dilute phase.

### Preparation of DNA structures for the benchmark systems

We first utilized the DNA structure building tool in GENESIS-CG-tool to generate a 200-bp double-stranded DNA (dsDNA) structure with a random sequence. Using the 3SPN.2C model^15, 16^, the 200-bp dsDNA system consists of 1,198 particles. Subsequently, we duplicated this dsDNA structure *n*^2^(*n* =1, 2, 3, 4, 5) times using GENESIS-CG-tool.

### Validation and benchmark simulations

For the benchmark tests, we first prepared droplet systems of TDP-43-LCD chains with four different particle numbers: (1) N_1_ =300,146, (2) *N*_2_ =744,128, (3) *N*_3_ =1,190,882, and (4) *N*_4_ =2,565,178. In all systems, there are two densities: *ρ*_*L*_ =6.74 ×10^−7^chains/Å^3^ and *ρ*_*H*_ = 1.85×10^−6^ chains/Å^3^.

Comparison tests between ATDYN (the atomistic decomposition method without dynamic load balancing), SPDYN-like (the mid-point cell method without dynamic load balancing), and CGDYN (the cell-based *kd*-tree domain decomposition with dynamic load balancing) are carried out for the systems with *N*_1_ particles on Fugaku with 4 MPIs per node. Performances are investigated by checking the wall time of 10,000 MD steps. We save the trajectory files at the final step to mimic the real MD simulations. In all cases, we run the same runs five times and get the average as the benchmark results. The effect of the load balancing update during MD simulations is checked for *N*_1_ particles with *ρ*_*H*_ density. In this case, we ran 10^8^ MD steps and saved trajectories at every 50,000 steps. In all cases, we assigned 35 Å as the electrostatic cutoff values (*r*_C,ele_ =35Å)

Benchmark comparison among ATDYN, CGDYN, and Open3SPN2 are carried out with the above-mentioned duplicated dsDNA systems. These systems have the cutoff distance *r*_c,ele_ =50 Å. For ATDYN and CGDYN, we used Intel Xeon Gold 6242 CPUs (32 cores per node). For the test with Open3SPN2, Nvidia RTX A6000 GPU cards are used. Note that we were unable to perform simulations of our *n*^2^ (*n* ≥ 2) dsDNA systems using the default Open3SPN2 software^40^.

We had to modify the structure-reading component of Open3SPN2 to make it functional. However, we did not make any changes to the kernel part responsible for force/energy calculations, so the performance should remain consistent with what was reported in the original paper^40^. In addition, we faced memory issues when utilizing Open3SPN2 for the duplicated dsDNA systems with *n*^2^ (*n* ≥ 6) 200-bp dsDNAs. However, we encountered no problems with such systems when using GENESIS ATDYN and CGDYN.

For the comparison between CGDYN and ATDYN for three systems on the RIKEN Hokusai supercomputer, we used the same working conditions as before^31^.

### CG MD simulations of droplet fusions

All CG simulations were performed using CGDYN. In all the simulations, we used a time-integration step size of 10 fs. Nonlocal HPS (*E*_HPS_) and electrostatic (*E*_*ele*_) potentials have cutoffs of 20 Å and 35 Å, respectively. Production runs were conducted in NVT ensembles, using Langevin dynamics with a friction coefficient of 0.01*P*s^-1^.

The two-droplet simulations were conducted in the periodic boundary condition (PBC) with box dimensions of 1000Å ×1000Å ×1500Å. MD simulations were carried out at 280 K for 5 ×10^8^ steps. However, we found that the fusion process occurred within a relative short time during the simulations. Therefore, only the first 1 ×10^8^ steps data are analyzed and shown in the current study. We performed the simulations on local PC clusters (Intel Xeon Gold 6240R CPU, 2.4GHz) using 16 MPI processes in conjunction with 3 OpenMP threads.

The ultra-large multiple-droplet simulations were conducted in PBC boxes of 2087Å ×2067Å ×2077Å (high-density) and 2899Å ×2903Å ×2929Å (low-density). We performed these simulations at *T* =290 *K* . The high-density and low-density systems were simulated for 4.25 ×10^8^ and 4.00 ×10^8^ steps, respectively. These simulations were conducted on supercomputer Fugaku using 512 or1,024 nodes.

## Data Analysis

The DBSCAN clustering method^42^ was employed to analyze the structures from the droplet simulations. However, different parameters were used for the two-droplet and multiple-droplet systems. In the two-droplet system, we defined the contact distance between two chains as *d*_*DBSCAN*,2−*drop*_=1 when the two chains formed contacts (i.e., when two inter-chain residues were within 10Å from each other), and 0 otherwise. On the other hand, in the multiple-droplet system, we define the contact distance between two chains as the distance between their COM. With these definitions, we used 𝜀 =0.5 (radius of neighborhood), min_pts=20 (the minimum number of neighbors for a point to be considered as a core point), and min_cluster_size=100 (minimum cluster size) in the two-droplet systems. For the multiple-droplet systems, we used 𝜀 =50Å, min_pts=5, and min_cluster_size=50. These parameters were chosen to effectively identify clusters and analyze the droplet structures in each system.

To monitor the mixing of contents in the two-droplet system, we define a coordinate *D*_*I,J*_ to describe the average distance between every two chains, one coming from cluster *I* and the other from cluster *J*:

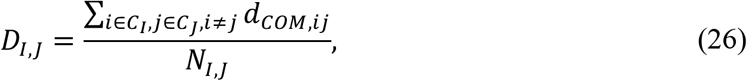

where *I* and *J* can be 1 or 2 (one of the two clusters in the initial structure), 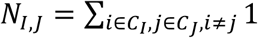 is the total number of computed chain pairs. On top of *N*_*I,J*_, we defined a “mixing” coordinate *m* as:

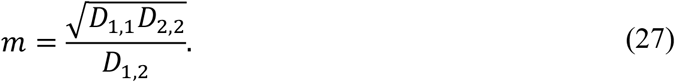

We employed a coordinate *η* to describe the shape of a droplet:

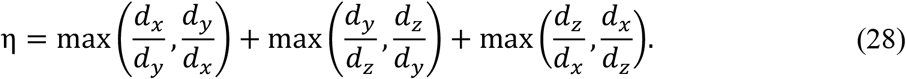

## Data availability

Benchmark and simulation data are available at https://github.com/RikenSugitaLab/cgdyntest/

## Code availability

All the codes for GENESIS CGDYN and the analysis programs are available at https://github.com/genesis-release-r-ccs/genesis-2.1.0beta_cgdyn

## Acknowledgements

This work was supported in part by MEXT JSPS Kakenhi (grant number 19H05645, 21H05249 (to Y.S.), 21H05282 (to J.J and C.T.)), RIKEN pioneering projects “Biology of Intracellular Environments”, and “Glycolipidologue Initiative” (to Y.S.), RIKEN incentive (to J.J), MEXT program for promoting research on the supercomputer Fugaku (JPMXP1020200101), and MEXT program for Big-data-driven bio/synthetic polymer science to create absolutely circular materials (JPMXP1122714694) (to Y.S.). The computer resources are provided by the HPCI system research project (Project ID: ra000003, hp200028, hp200135, hp210177, hp220170, hp230072, and hp230212) and by RIKEN Advanced Center for Computing and Communication (for HOKUSAI BigWaterfall, project Q22535, Q22536, and Q23536).

---

**Supporting Figure 1.**
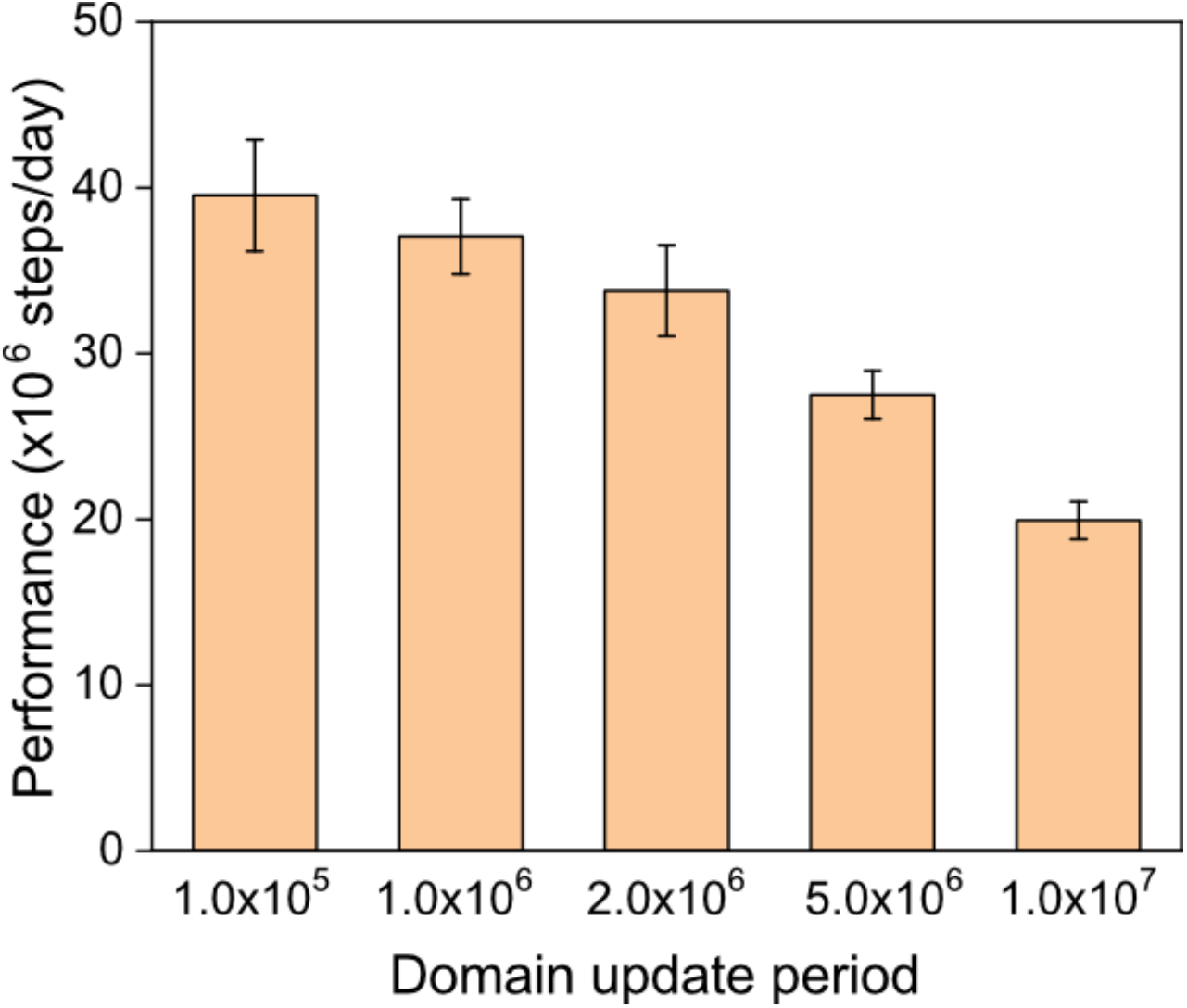
Performance of droplet systems by changing the domain update period.

**Supporting Figure 2.**
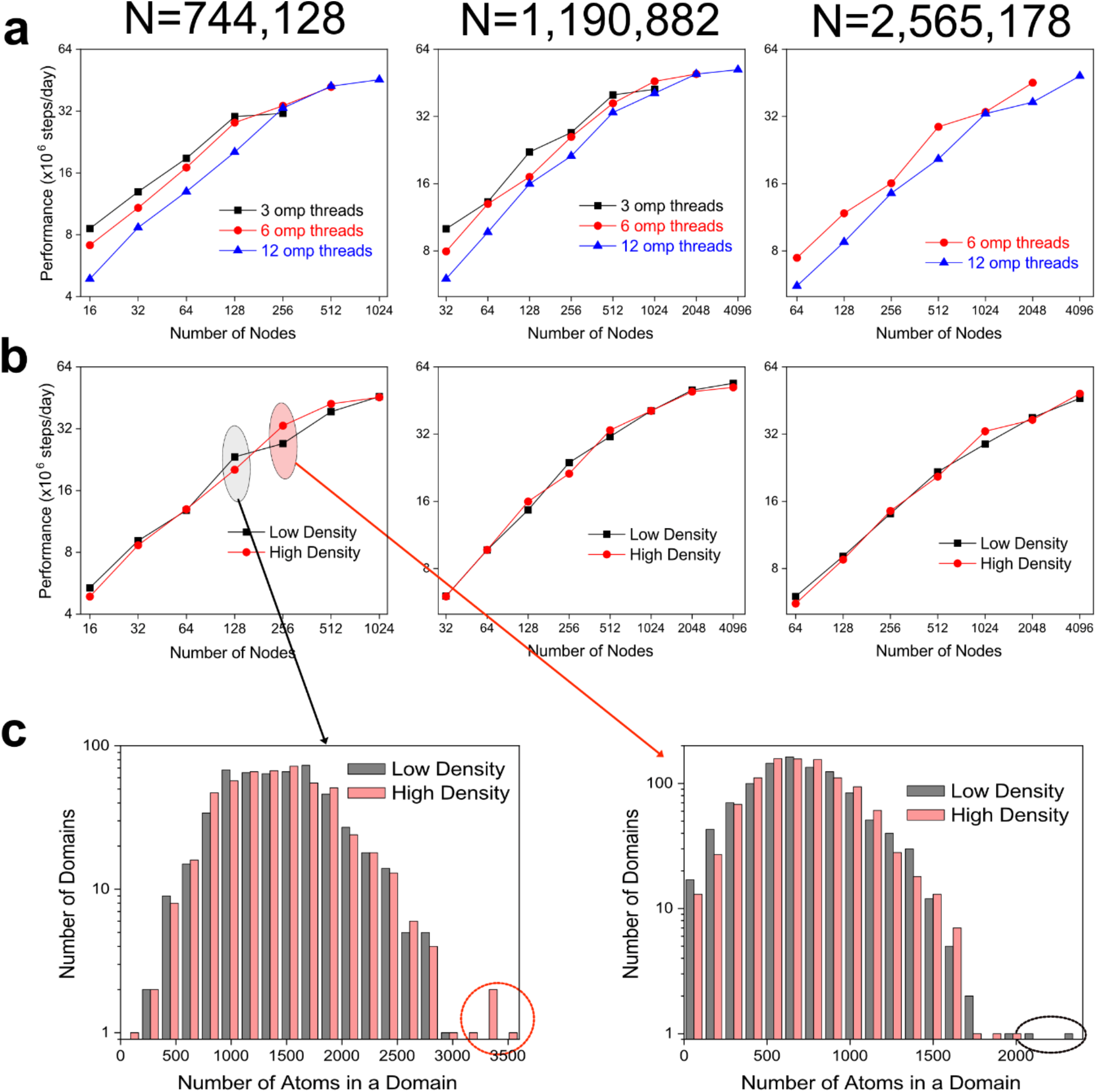
Benchmark results of droplet systems in CG MD simulations with CGDYN. (a) Benchmark performance of a high-density system as a function of the number of nodes, changing the number of OpenMP threads. (b) Comparison of the performances between the high- and low-density systems. There is no significant performance difference between them. (c) The distributions of the number of particles in domains with CGDYN. The performance is limited by the maximum number of particles in a domain.

**Supporting Figure 3.**
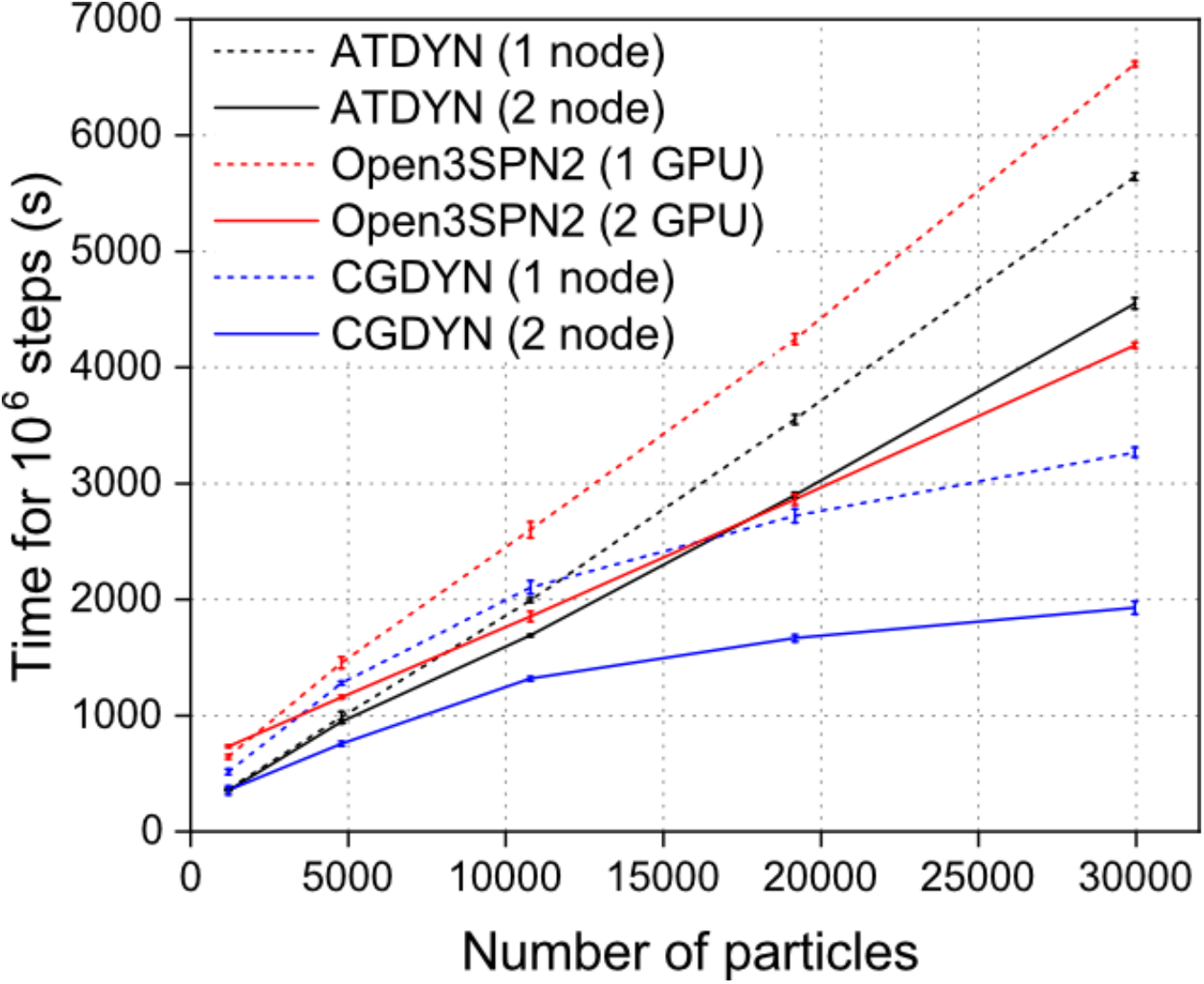
Comparison of the performance of multiple DNA chains. For a very small system, ATDYN shows the best performance. As the system size increases, CGDYN shows better performance than ATDYN and Open3SPN2. CGDYN has the best performance irrespective of the system size, just using 2 nodes.

**Supporting Figure 4.**
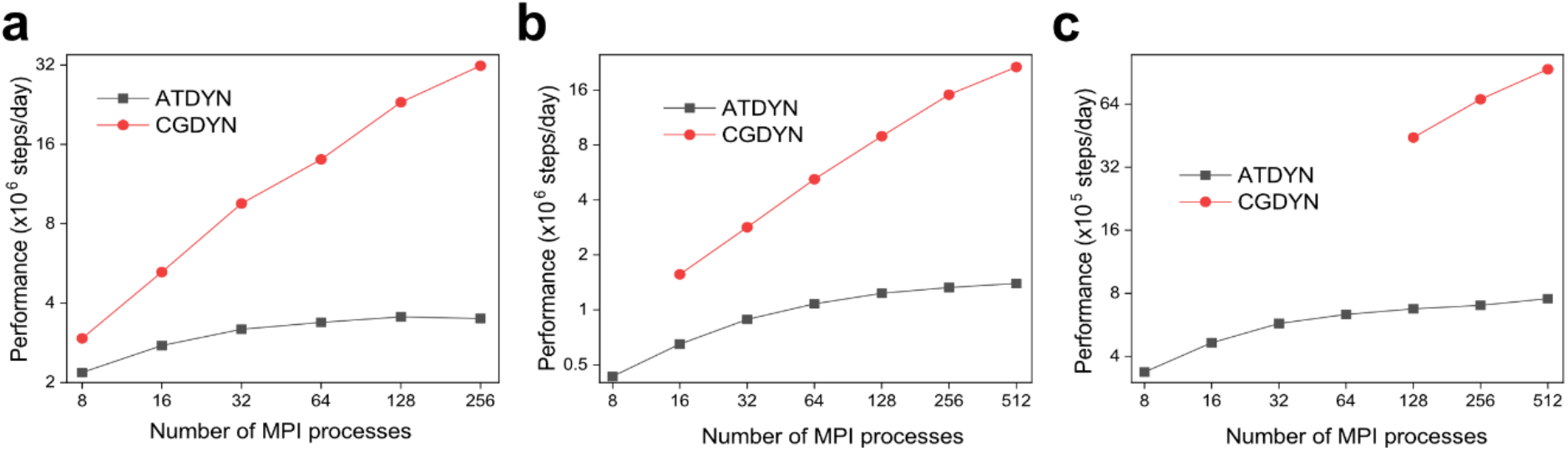
Performance of MD simulations using CGDYN for (a) 120 DPS proteins (222,360 particles), (b) 5,000 chains of 100 amino acid IDPs (500,000 particles), and (c) 512 nucleosomes (1,044,480 particles) on the RIKEN Hokusai supercomputer.

**Supporting Figure 5.**
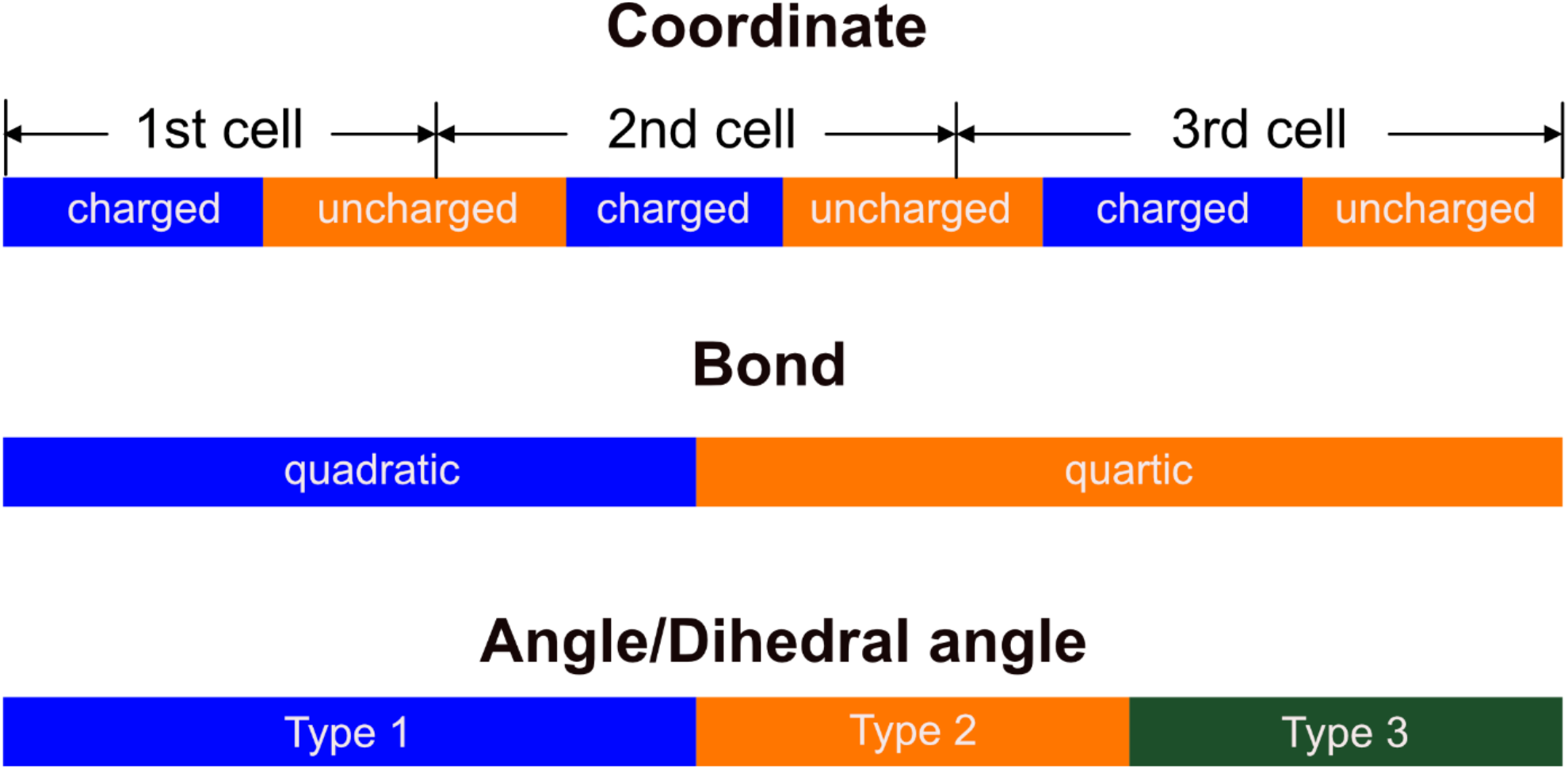
Array structures of the coordinates, bonds, and angles/dihedral angles. In each cell, we save all the charged and uncharged particles in this order. Bond/Angle/Dihedral angle and other bonded interaction data are not saved cell-wise. There are multiple potential functions for the same bonded term. We save the same potential function terms consecutively.

**Supporting Figure 6.**
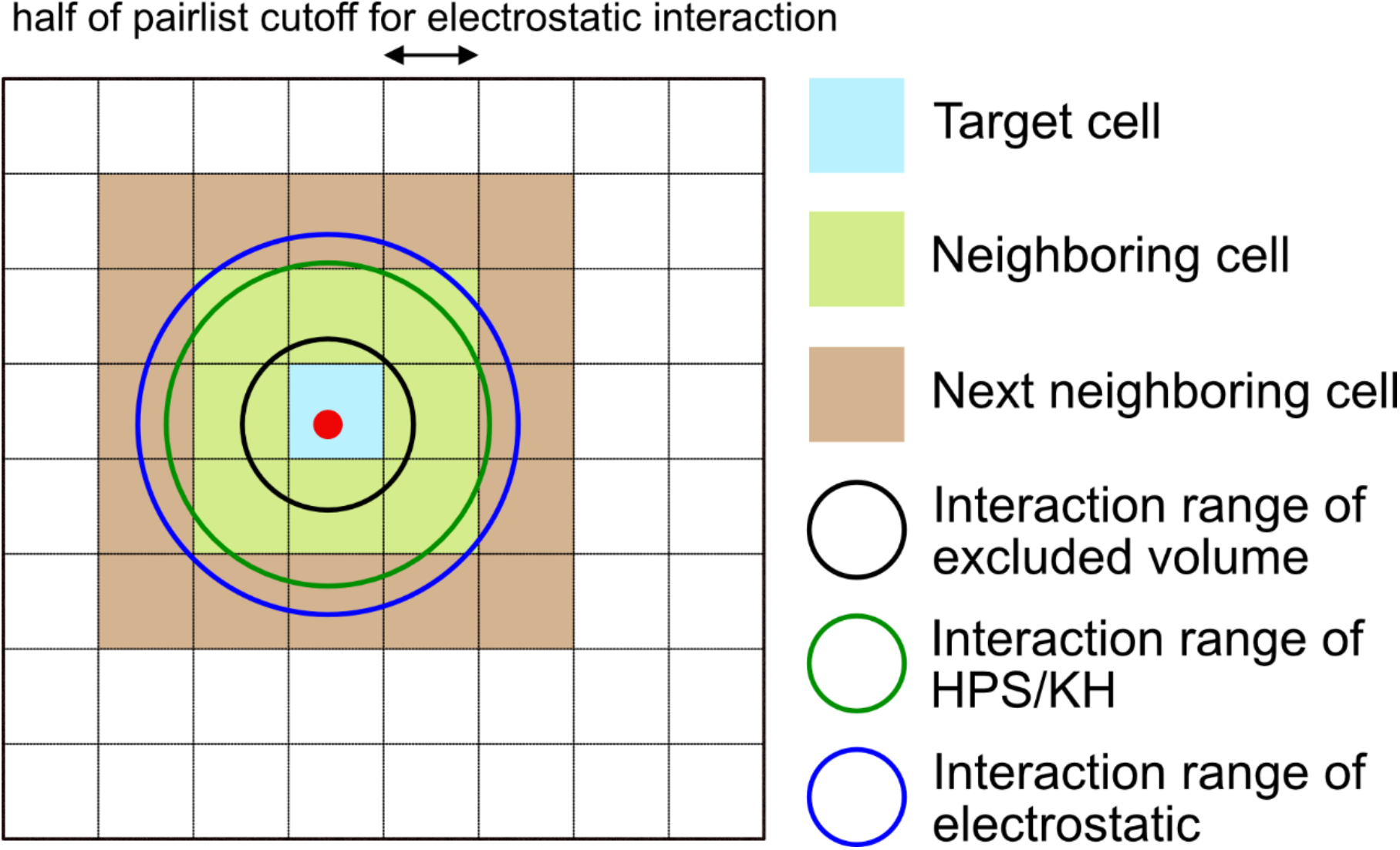
Interaction ranges of potential functions. For a particle marked as a red circle, the interaction ranges of the excluded volume, HPS, and electrostatic potentials are represented as black, green and blue circles, respectively. Because of the short interaction range of the excluded volume, we can consider only particles in the cells, including the particles in the target cell and those in the neighboring cells. The interaction range of HPS/KH and electrostatic is much longer, and we consider the particles in the next neighboring cells as well as the target/neighboring cells.

**Supporting Figure 7.**
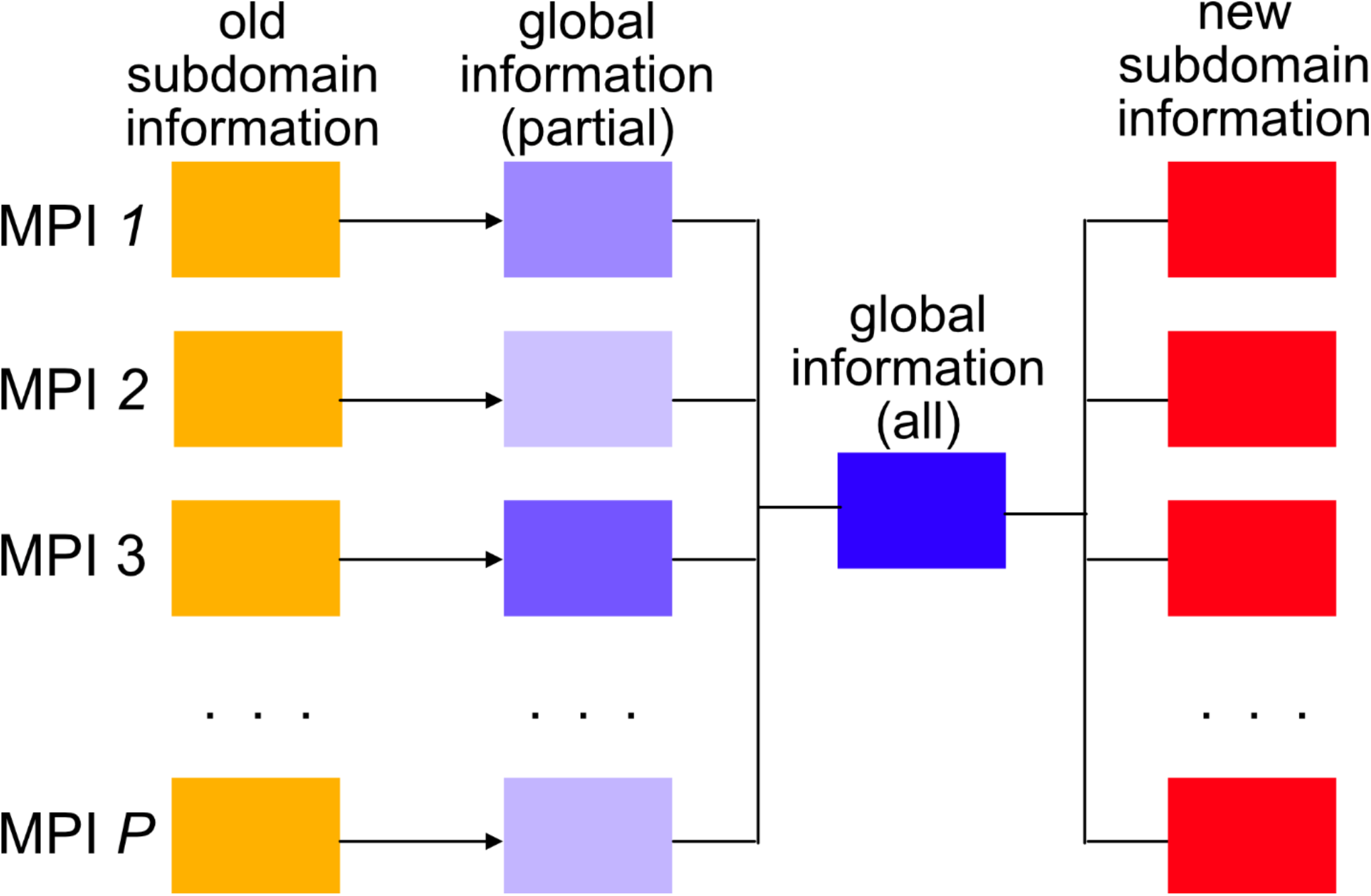
Cell reassignment in dynamic load balancing. First, each process saves its subdomain data into a global information array. After collective communications, all processes share the global information data of all particles and redefine subdomains based on the particle densities from the global information data.

## Supporting Algorithm. Neighbor List generation in CGDYN

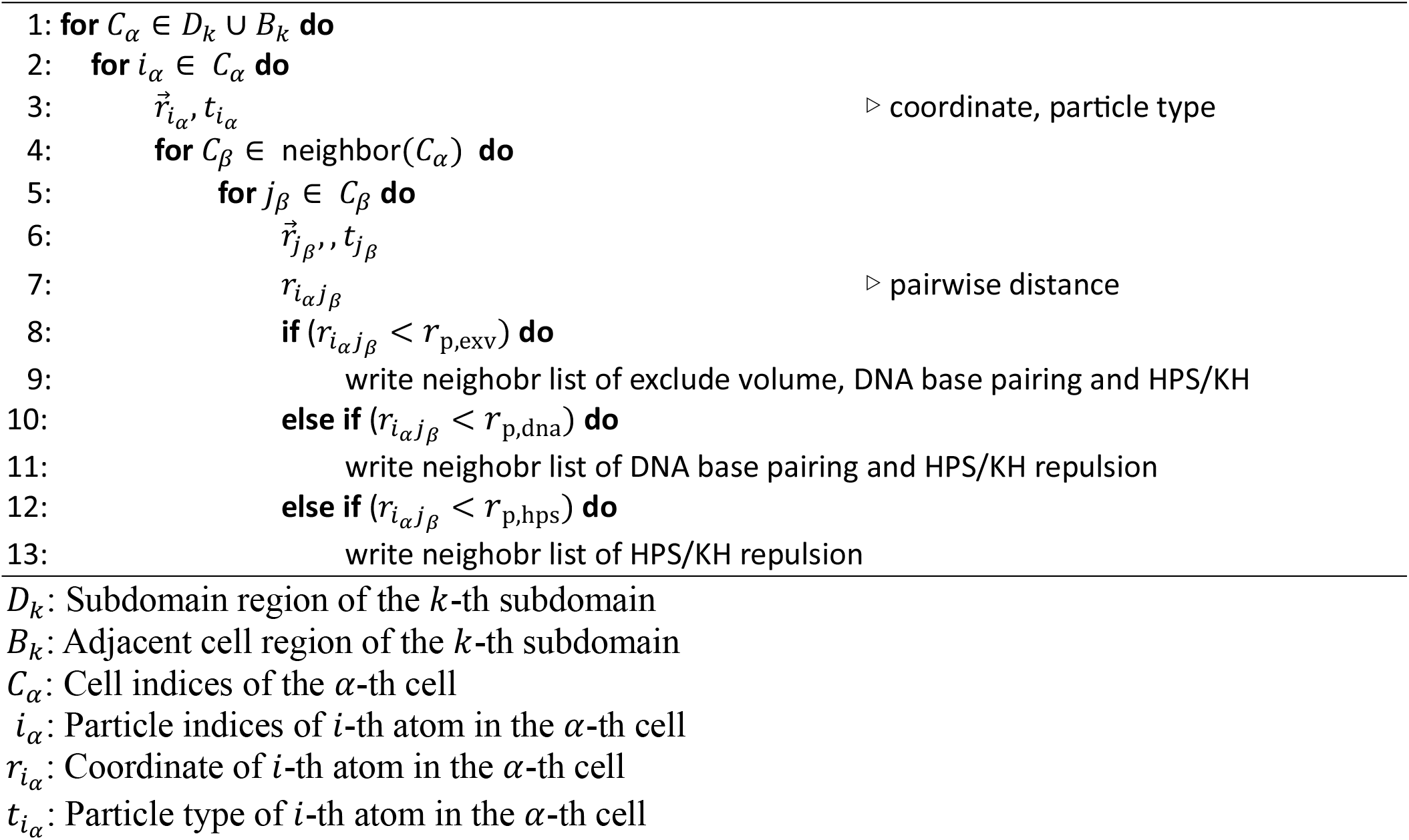

**Supporting Table 1.** Benchmark performance of multiple DNAs (unit is 10^6^ steps/day).

| Number of Nodes | Number of particles = 119,800 |  | Number of particles = 269,550 |  | Number of particles = 479,200 |  |
| --- | --- | --- | --- | --- | --- | --- |
|  | ATDYN | CGDYN | ATDYN | CGDYN | ATDYN | CGDYN |
| 1 | 3.576 |  | 1.475 |  | 0.807 |  |
| 2 | 4.557 |  | 1.916 |  | 1.056 |  |
| 4 | 5.437 |  | 2.294 |  | 1.247 |  |
| 8 | 3.576 | 15.393 | 2.509 | 10.363 | 1.380 |  |
| 16 | 6.278 | 20.433 | 2.728 | 17.728 | 1.520 | 9.876 |
| 32 | 5.922 | 29.993 | 2.739 | 27.078 | 1.560 | 13.789 |
| 64 | 5.548 | 42.667 | 2.743 | 42.562 | 1.617 | 20.241 |
| 128 | 4.192 | 44.158 | 2.494 | 40.393 | 1.506 | 25.681 |

